# Neural encoding of the reliability of directional information during the preparation of targeted movements

**DOI:** 10.1101/2020.09.17.302000

**Authors:** Charidimos Tzagarakis, Sarah West, Giuseppe Pellizzer

## Abstract

Visual information about the location of an upcoming target can be used to prepare an appropriate motor response and reduce its reaction time. Here, we investigated the brain mechanisms associated with the reliability of directional information used for motor preparation. We recorded brain activity using magnetoencephalography (MEG) during a delayed reaching task in which a visual cue provided valid information about the location of the upcoming target with 50, 75 or 100% reliability. We found that reaction time increased as cue reliability decreased and that trials with invalid cues had longer reaction times than trials with valid cues. MEG channel analysis showed that during the late cue period the power of the beta-band from left mid-anterior channels, contralateral to the responding hand, correlated with the reliability of the cue. This effect was source localized over a large motor-related cortical and subcortical network. In addition, during invalid-cue trials there was a phasic increase of theta-band power following target onset from left posterior channels, localized to the left occipito-parietal cortex. Furthermore, the theta-beta cross-frequency coupling between left mid-occipital and motor cortex transiently increased before responses to invalid-cue trials. In conclusion, beta-band power in motor-related areas reflected the reliability of directional information used during motor preparation, whereas phasic theta-band activity may have signaled whether the target was at the expected location or not. These results elucidate mechanisms of interaction between attentional and motor processes.

## Introduction

A key function of cognition is the integration of information for predictive processing (Bubic et al., 2010). In particular, in time-stressed tasks such as, for example, while playing tennis or car driving, the response latency decreases when the motor response is correctly anticipated (Shim et al., 2005; Stahl et al., 2014). However, if the anticipation is incorrect and the response planned needs to change, then the latency is lengthened (Posner et al., 1980). Thus, the reliability of the information used to anticipate the response plays an important role in the effectiveness of motor preparation (Vossel et al., 2014; Arjona et al., 2016). Studies of spatial attention using visually cued tasks have shown that the greater the proportion of valid-cue trials within a block of trials, that is, the greater the reliability of the information the cue conveys, the greater the reduction in response latency to correctly anticipated targets (Jonides, 1980; Posner et al., 1980; Eriksen and Yeh, 1985; Risko and Stolz, 2010; Arjona et al., 2016; Kuhns et al., 2017; Valakos et al., 2020). In this context, motor planning and spatial attention are inherently linked (Goldberg and Segraves, 1987; Rizzolatti et al., 1987; Rushworth et al., 2003; Brown et al., 2011; Perfetti et al., 2011), and several studies have shown that the frontoparietal neural network associated with motor preparation overlaps to a large extent with the network associated with spatial attention (Goldberg and Segraves, 1987; Corbetta et al., 1998; Nobre et al., 2000; Moore and Fallah, 2001; Rushworth et al., 2003; Balser et al., 2014; Denis et al., 2017). Therefore, we can expect that changing the reliability of information provided by visual cues regarding an upcoming motor response will be reflected in changes of neural activity in the frontoparietal networks associated with motor control and spatial attention.

It is well known that during motor preparation, there is a reduction in power of beta-band (15-30 Hz) oscillations over the peri-Rolandic region (Jasper and Penfield, 1949; Pfurtscheller, 1981) to a level that is intermediate between baseline and the even lower level associated with motor execution. However, how much beta power decreases before movement onset varies with the degree of motor preparation (Alegre et al., 2003; Tzagarakis et al., 2010; Grent-’t-Jong et al., 2015; Tzagarakis et al., 2015; Tzagarakis et al., 2019) and is reflected by a corresponding change in response latency and number of premature responses (Tzagarakis et al., 2010; Tzagarakis et al., 2015; Tzagarakis et al., 2019; Barth et al., 2021). Furthermore, it was shown in motor choice tasks that sensorimotor beta power continuously reflects the probabilistic inference based on the accumulation of evidence in favor of one or another motor response (Donner et al., 2009; Gould et al., 2012). For these reasons, we hypothesized that the reliability of the information provided by visual cues modulates the power of beta-band activity in motor-related areas during motor preparation.

To investigate the brain mechanisms associated with the reliability of visual information about the location of an upcoming target, we recorded whole-head neuromagnetic activity using magnetoencephalography (MEG) during a visually cued reaching task in which the reliability of the cue varied across blocks of trials. We did not limit our analysis to the beta-band, but also checked whether there was any unanticipated effect of cue reliability on other frequency bands (delta, theta, alpha, and gamma). We analyzed the neural data during the early cue period associated with the phasic change of activity following the onset of the cue, and during the late cue period which is associated with a more stable pattern of activity. In addition, since cue reliability was defined by the proportion of cue-valid trials within a block of trials, we checked whether there was a change in general arousal level associated with the reliability condition during the baseline period.

In addition, we tested the neural activity after target onset, that is, once the validity of the cue was determined. Previous studies have shown that theta-band (4-7 Hz) activity plays an important role in signaling an unanticipated visual stimulus (Rawle et al., 2012; Proskovec et al., 2018). In particular, Rawle et al. (2012), using a visually cued reaching task, found that there was a parietal and frontal phasic increase in theta-band power after target onset when the location of the target was uncertain but not when it was known in advance. Consequently, in visually cued reaching tasks, we expected the occurrence of a change in theta-band power in frontoparietal areas after target onset of invalid-cue trials, that is, when the cue does not provide valid information about the location of the upcoming target. However, here too we examined whether there was an unanticipated effect of cue validity on the other frequency-bands.

Finally, we investigated the interaction between attentional and motor processes by analyzing the functional connectivity of the brain regions found to be associated with cue reliability and validity. From the hypotheses mentioned above, we expected to find different theta-beta cross-frequency coupling depending on the validity of the cue.

## Materials and Methods

### Participants

Twelve right-handed volunteers with normal or corrected-to-normal vision participated in this study (7 women and 5 men; mean age=31 years; age range 24-45 years). Participants had no reported history of neurological or psychiatric disorders, no history of substance abuse, and no tobacco use for at least one month prior to the recording session. They provided written informed consent prior to being included in the study and received monetary compensation for their participation. The protocol was approved by the Institutional Review Board of the Minneapolis Veterans Affairs Health Care System.

### Task setup

Participants used their right hand to control the position of a cursor on a screen using a joystick (model M11C0A9F customized for MEG compatibility, CH Products, Vista CA, USA). A trial was initiated by placing the cursor within a small circular window (diameter=1 deg of visual angle) in the center of the screen for a 3 s center-hold period, and by fixating the center. After the center-hold period, the cue (empty circle; diameter=2 deg of visual angle) was presented at 4 deg of visual angle from the center of the screen. The direction of the cue varied randomly from trial to trial in the 0-360 deg range. The duration of the cue period was selected randomly from a uniform distribution over the 1.0-1.5 s interval to reduce the effect of temporal expectancy (Luce, 1986; Wagener and Hoffmann, 2010). Participants were instructed to fixate the center of the screen during the center-hold and cue periods. Note that central fixation was not required after target onset to facilitate the task, however, given the size of the target and its eccentricity well within parafoveal vision, the task could be performed without moving the eyes. The direction of gaze was monitored on-line using a video-based eye tracking system (ISCAN ETL-400, ISCAN Inc., Woburn, MA). If the subjects blinked or did not maintain fixation within 2 deg from the center, the trial was automatically aborted, and a new trial had to be initiated. For three participants, the fixation threshold had to be increased during the second and/or third block of trials due to a drift of gaze calibration. In addition, the artifact rejection process (described further below) was used to reject any trial contaminated by eye movements that might not have been detected during the recording. After the cue period, the target (filled circle) appeared at the position of the cue in case of a valid-cue trial, or at a different random direction around the center in case of an invalid-cue trial. Participants were instructed to move the cursor quickly and accurately from the center onto the target. A schematic representation of the task is shown in Figure 1A. The reaction time (RT) was defined as the interval between the onset of the target and the exit of the cursor from the center window. RTs shorter than 100 ms or longer than 1,500 ms were counted as errors. The movement time (MT) was defined as the interval between when the cursor exited from the center window to when it entered the target. MTs greater than 1,500 ms were considered errors. For a successful movement execution, the trajectory of the cursor had to remain within virtual boundaries tangent to the center window and the target, and the cursor had to remain on the target for a minimum of 100 ms. If any of these conditions did not hold, the trial was aborted and presented again at a random position in the sequence of the remaining trials in the block. Consequently, every condition had the same number of correct trials. At the end of the trial, an auditory signal indicating whether the trial was correct (high pitch) or not (low pitch) provided feedback to the participant. An interval of at least 3 s separated each trial. All participants practiced the task before the MEG recording.

**Figure 1.**
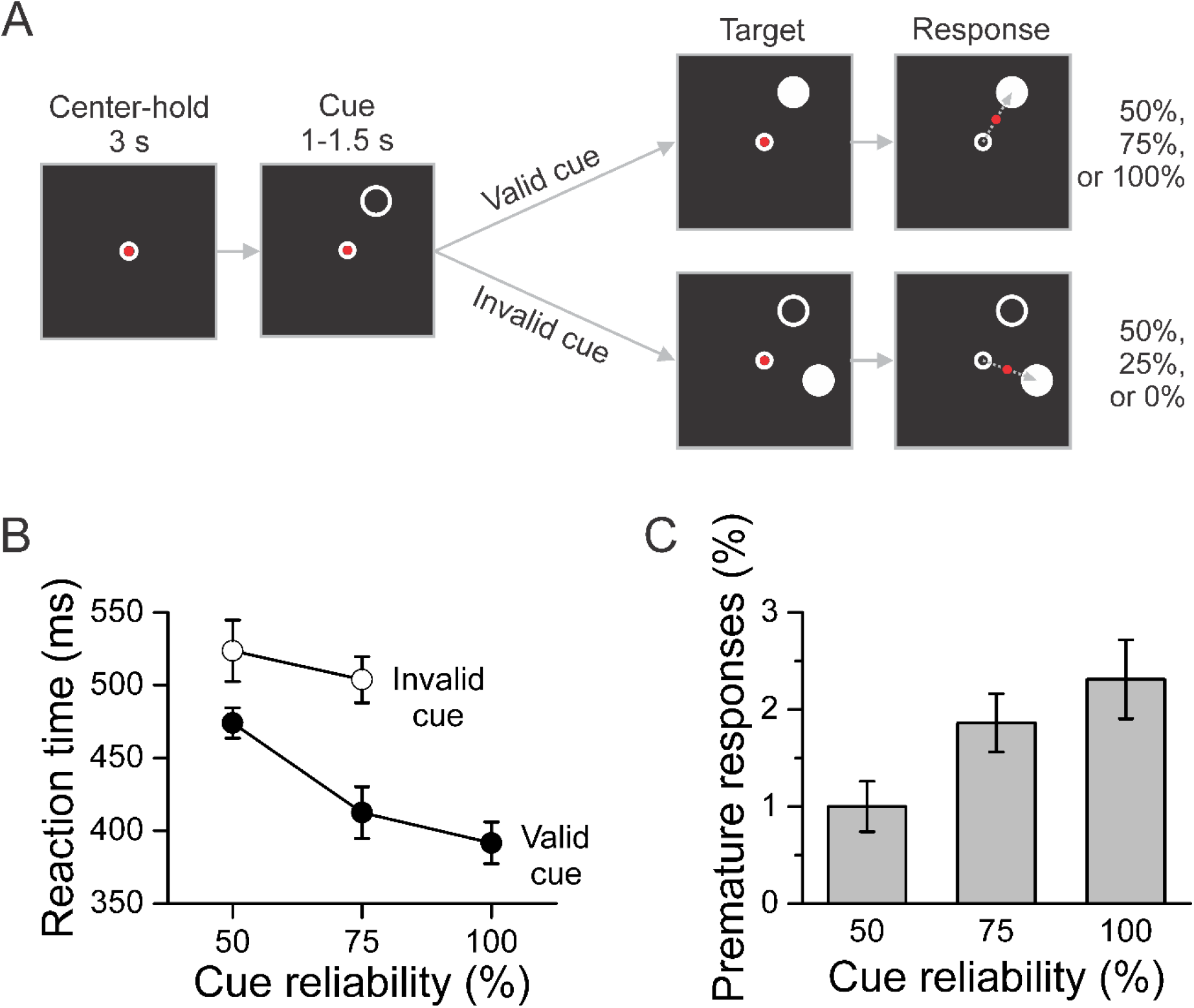
Task and behavioral results. ***A,*** Schematic description of the task. After a 3 s center-hold period, a peripheral cue (white circle) was presented for 1 to 1.5 s in any direction around the center. The direction of the cue was selected randomly in the 0-360 deg range from trial to trial. The cue indicated the location of the upcoming target (white disc) with 50%, 75%, or 100% reliability depending on the block of trials. Participants were informed of the cue reliability condition before the start of the block of trials. After target onset, which determined whether the trial had a valid or invalid cue, participants responded by moving a joystick-controlled cursor (red dot) from the center of the screen to the target as quickly and accurately as possible. ***B,*** Mean reaction time across participants per cue reliability and cue validity conditions. Mean reaction time decreased significantly as cue reliability increased and was significantly shorter for valid-cue trials than for invalid-cue trials. ***C,*** Percent of premature responses per cue reliability condition averaged across participants. The percent of premature responses increased significantly with cue reliability. The error bars indicate the SEM (N=12 participants) for within-subject designs (Cousineau, 2005; Morey, 2008).

### MEG recordings

Data acquisition was performed similarly to previously reported experiments (Tzagarakis et al., 2010; Tzagarakis et al., 2015). Participants were lying supine on a bed inside of a magnetically shielded room with their head in the MEG detector helmet. The visual stimuli and joystick-controlled cursor were projected on a screen about 60 cm in front of the subject using an LCD video projector (Sony VPL-PX20) located outside of the shielded room. The joystick was secured to the bed next to the subject’s right hip so that it could be manipulated comfortably with the right hand. Neuromagnetic signals were recorded using a 248-channel whole-head MEG system equipped with first-order axial gradiometers (Magnes 3600 WH, 4D Neuroimaging, San Diego CA, USA). The signals were low-pass filtered (DC-400 Hz) and sampled at a rate of 1017.25 Hz. An electrooculogram (EOG) was recorded in addition to the video-based eye-tracking signal to help identify epochs contaminated by eye movements or eye blinks. In addition, the onset of the visual stimuli (cue and target) was detected by a photodiode to ensure timing accuracy. The video-based eye-tracking, EOG, photodiode and joystick signals were recorded on auxiliary channels of the MEG system to ensure synchronization of all the data. Five small coils were attached on the subject’s head to measure the position of the head relative to the gradiometers at the beginning and end of the recording session. The head shape of each subject was digitized using a 3-D digitizer (Fastrak, Polhemus, Colchester VT, USA). In addition, the position of three fiducial points (nasion, left and right pre-auricular points) was also digitized. MEG data were analyzed using the open-source toolbox Fieldtrip (Oostenveld et al., 2011) and MATLAB (Mathworks Inc., Natick MA) custom-written programs. Signals from reference gradiometers were used to remove background noise from the neuromagnetic data using the 4D-Neuroimaging algorithm implemented in Fieldtrip. Data were aligned to either the onset of the cue or the onset of the target, and trials contaminated by electronic artifacts, eye movements, eye blinks, or muscle activity were detected using a data-adaptive threshold and discarded. Following the artefact rejection procedure, 92% (95% CI: 89%-95%) of cue-aligned trials and 91% (95% CI: 87%-94%) of target-aligned trials were kept for analysis. There was no significant difference in the proportion of trials kept for analysis across cue reliability conditions (cue-aligned: F(2,33)=1.865, p=0.171; target-aligned: F(2,33)=1.724, p=0.194). Independent component analysis was applied to identify components with cardiac artifacts, which then were removed from the data. Finally, the data were detrended and an adaptive anti-aliasing low-pass filter was applied before resampling at 256 Hz to facilitate further processing. For each participant, one MEG channel was malfunctioning and removed from all analyses.

### MRI

Head anatomical magnetic resonance images (MRI) were obtained to help estimate the sources of the MEG channel-level results. T1-weighted images were acquired with a 3-dimensional multiplanar gradient echo sequence using a 3 Tesla system (Achieva, Philips Medical Systems, Andover MA, USA; repetition time=8.0744 ms; echo time=3.695 ms; flip angle=8°; field of view=240×240 mm; matrix=256×256 pixels; slice thickness=1 mm). The volume of the MRI anatomical scan extended from the top of the head to the bottom of the cerebellum and included all fiducial points. MRIs were obtained from ten out of twelve participants. Two of the participants did not have an MRI and were associated with a set of images from another participant with a similar head size and shape.

### Experimental Design and Statistical Analyses

#### Task

Participants performed an instructed-delay reaching task in which a peripheral visual cue indicated the location of the upcoming target with 50%, 75% or 100% reliability depending on the block of trials. That is, the cue was either valid when the target was presented at the same location as the cue, or invalid when the target was presented at a different location. Consequently, each trial was defined by the reliability condition (50%, 75%, 100%) of the block of trials, and by the validity (valid, invalid) of the cue. Participants were informed verbally about the reliability condition before each block of trials, and each block was composed of 72 trials. The order of the reliability conditions was fully counterbalanced across participants.

#### Behavioral data analysis

For each participant, the average RT and MT per cue reliability and cue validity condition was computed using the harmonic mean which is robust to potential outliers (Ratcliff, 1993). Several competing GLMM were tested and those with the best Akaike information criterion were selected (McCulloch et al., 2008). First, we verified whether decision time as measured by reaction time was stable across trials within each block of reliability conditions. To this end, for each block of trials we formed twelve consecutive sub-blocks of six correct trials and computed mean RT for each sub-block. We tested whether the block order (i.e., order of reliability conditions), the sub-block of trials, or their interaction had a significant effect on mean RT using a GLMM with a gamma distribution, a log link function, and an AR1 covariance structure. For the main analysis, RT and MT were analyzed using a GLMM with a gamma distribution and log link function, whereas the proportion of errors was analyzed using a GLMM with the binomial distribution and probit link function. Participants were treated as a random effect. The effects of cue reliability, cue validity, and their interaction were tested on RT, MT and proportion of directional errors, whereas only the effect of cue reliability was tested for errors preceding target onset (i.e., fixation errors and early response errors) since cue validity was not yet determined at that point in the trial. These analyses were implemented using SPSS (version 24; IBM corp., Armonk, NY).

Given the size of the target (2 deg of visual angle) and its eccentricity (4 deg of visual angle), the task could be performed without moving the direction of gaze toward the target. However, to facilitate the task, we did not enforce central fixation after target onset (see Task Setup, above). For that reason, we analyzed the oculomotor activity from target onset until when the cursor reached the target. To this end, we computed the mean and SD of the X and Y coordinates of eye position during 250 ms of central fixation before target onset. We used 3 SD as the threshold to detect eye movements that were made outside of central fixation.

#### MEG channel-level analysis

We analyzed the effect of the experimental factors on the power of the delta (1-3 Hz), theta (4-7 Hz), alpha (8-12 Hz), beta (15-30 Hz) and gamma (50-80 Hz) bands. Time-varying power in each band was obtained after applying Finite Impulse Response (FIR) bidirectional filters to the broadband MEG signal. To test whether there was a significant effect of cue reliability on band power, we computed the dependent samples regression between log-transformed power and cue-reliability for each frequency-band and for three time windows of interest: (1) baseline (0.6-0.1s before cue onset), (2) early cue period (0-0.5 s after cue onset), and (3) late cue period (0.5-1.0 s after cue onset). The statistical significance of the regression was determined using a nonparametric cluster-based test with a permutation distribution of 10,000 rearrangements of data from the three reliability conditions within twelve subjects (Maris and Oostenveld, 2007; Tzagarakis et al., 2015). The significance threshold was set at p<0.0033 after using Bonferroni correction for testing five frequency bands and three time-windows and keep the family-wise error rate at p<0.05. Note that since we were interested to check for a potential effect of cue reliability on the baseline period, we did not compute power relative to baseline. However, the statistics extracted and the methods used to compute the p-values are independent from differences in baseline across subjects. Furthermore, the SEM represented in the figures were computed using the Cousineau-Morey method for within-subject designs which controls for between-subjects variance (Cousineau, 2005; Morey, 2008). In addition, we tested the effect of cue validity on the power of each frequency band during the target presentation period (0-0.4 s after target onset) using a paired *t*-test between the log-transformed power during valid-cue and invalid-cue trials. The statistical significance was obtained using the same cluster-based permutation approach described above with the distribution of all 4,096 permutations of two validity conditions within twelve subjects. The significance threshold level of each analysis was set at p<0.01 after using Bonferroni correction for testing five frequency bands.

#### Source-level analysis

The head MRI of each subject were segmented in order to create a single-shell model of the brain surface (Nolte, 2003). The brain volume was divided in a regular grid of 6 mm isotropic voxels which was normalized into Montreal Neurological Institute (MNI) brain space (Evans et al., 2012). The co-registration between MRI and MEG data was performed using the three fiducial points and the head shape. The lead field matrix was computed for each grid location. The localization of neural sources was performed using the Dynamic Imaging of Coherent Sources (DICS) beamformer using a regularization parameter of 10% (Gross et al., 2001). Based on the channel-level results, we performed source localization analysis for the correlation of beta-band power with cue reliability during the late cue period. In addition, source estimates for beta-band power during the reaction time and the baseline period were obtained for the purpose of the theta-beta cross-frequency coupling analysis (see below). In each case we used trials from all cue reliability conditions to compute the common spatial filter using multi-taper Fourier transform for the cross-spectral density estimation. We also performed source analysis for the effect of cue validity on theta-band power during the reaction time period. In this case, we used all the valid-cue and invalid-cue trials from the 50% and 75% conditions to compute the common spatial filter, and the cross-spectral density was computed from a Fourier transform using a Hanning taper. The common spatial filters were used to estimate the beta-band source activity during the late cue period for each reliability condition, and to estimate the theta-band activity after target onset for valid-cue and invalid-cue trials of the 50% and 75% reliability conditions. Voxelwise dependent samples regressions for log-transformed power across reliability conditions, and paired *t*-tests for valid-cue and invalid-cue trials were calculated as described for the channel-level analysis. Finally, groups of voxels were selected using the same nonparametric cluster-based permutation method as for the channel-level analysis (Maris and Oostenveld, 2007).

#### Theta-beta cross-frequency coupling

Since we found a significant effect of cue validity on theta power during the reaction time (see Results), and since it is well-known that there is a strong reduction of beta power during the reaction time (Pfurtscheller, 1981; Tzagarakis et al., 2010; Tzagarakis et al., 2015; Heinrichs-Graham and Wilson, 2016), we investigated whether there was a theta-beta cross-frequency interaction during that period. To this end, we computed the pairwise phase consistency (PPC), which is an unbiased and consistent estimator of rhythmic synchronization (Cohen, 2008; Vinck et al., 2010; Aydore et al., 2013). PPC was computed between the phase of the theta-band signal for each voxel selected by the cluster-based analysis of the effect of cue validity (Fig. 4 and Fig. 5A, yellow) and the amplitude of the beta-band of each voxel in the left precentral gyrus that was selected by the cluster-based analysis of the decrease in beta power from baseline during the 0-0.4 s period after target onset (Fig. 5A, blue). The average PPC per voxel for the 0.1-0.3 s following target onset was compared between the valid-cue and invalid-cue conditions with a paired *t*-test followed by a cluster-based analysis (Maris and Oostenveld, 2007). The procedure was implemented as follows. The signal at each voxel analyzed was computed using the Linearly Constrained Minimum Variance (LCMV) beamformer (Van Veen et al., 1997), using the broadband signal data covariance matrix and a regularization parameter of 10%. The LCMV beamformer was based on the cross-correlation matrix estimated from all valid-cue and invalid-cue trials from the 50% and 75% cue reliability conditions. The broadband signal for each trial and voxel was obtained by multiplying the channel space signals with the beamformer solution. Source signals for each trial were then band-pass filtered at the frequency band of interest using a bidirectional Finite Impulse Response (FIR) filter. The time-series of the phase of the theta-band and of the amplitude of the beta-band were computed from the analytical signal obtained using the Hilbert transform. The time-series of the phase of the beta-band amplitude was calculated after de-meaning the data (Cohen, 2008). PPC values from each theta-band voxel to each of the beta-band precentral cortex voxels were then averaged across conditions (50 and 75 % reliability). This resulted in a dataset of PPC time-series for each theta-band voxel in each of the two cue-validity conditions. We tested the effect of cue-validity on theta-beta PPC during the 0.1 to 0.3 s period following target-onset using a paired *t*-test between the average PPC of valid-cue- vs. invalid-cue trials. Clusters of voxels were selected using the same nonparametric cluster-based permutation test mentioned above (Maris and Oostenveld, 2007). Furthermore, for each of the two cue-validity conditions, we extracted the PPC time-series for each selected voxel for further analysis.

#### Potential pitfalls of the PPC analysis

Connectivity metrics, such as the PPC, are susceptible to several potential pitfalls (Palva and Palva, 2012; Aru et al., 2015; Bastos and Schoffelen, 2015). Prominent amid these is the problem of field-spread which causes activity originating from a single source being captured by several sensors. However, this potential confound is mitigated by performing the analysis in source space and by comparing the same subjects in different conditions (Schoffelen and Gross, 2009; Palva and Palva, 2012). We also addressed the possibility that a difference in PPC (i.e., ΔPPC) between valid-cue and invalid-cue conditions resulted from the difference in theta-band power between conditions (Schoffelen and Gross, 2009; Aru et al., 2015; Bastos and Schoffelen, 2015). To this end, we created a bootstrap distribution of *i*=1,…1,200 ΔPPC_*i*_ averaged across subjects as follows. For each re-sampling iteration, the target-aligned time-series of the phase of the theta-band was kept as in the original dataset, whereas the time-series of the amplitude of the beta-band was extracted from the corresponding trial but starting at a random time during the trial. If ΔPPC in the original dataset was dependent on the power of theta, then the same effect would occur with the re-sampled dataset as with the original dataset.

## Results

### Behavioral performance

#### Reaction time and movement time

We found no significant change of mean RT due to block order (F(2,396)=0.556, p=0.574), sub-block of trials (F(11,396)=1.319, p=0.211), or their interaction (F(22,396)=0.997, p=0.467). Consequently, this analysis indicates that there was no significant change of mean RT across the succession of blocks and sub-blocks of trials. For this reason, all trials have been used for the subsequent behavioral and MEG data analyses.

Mean RT per cue reliability and cue validity is plotted in Figure 1B. The GLMM analysis showed that, as expected, it decreased significantly as cue reliability increased (F(2,55)=15.706, p<0.001), and that it was significantly shorter for valid-cue trials than for invalid-cue trials (F(1,55)=18.427, p<0.001). There was no significant reliability x validity interaction (F(1,55)=2.159, p=0.147). In contrast to RT, we found no significant effect of cue reliability, cue validity or their interaction on MT (reliability: F(2,55)=2.155, p=0.126; validity: F(1,55)=0.040, p=0.843; interaction reliability x validity: F(1,55)=0.358, p=0.552). Mean MT across subjects was 188.3 ms (SEM=26.7 ms, N=12).

#### Error trials

The implementation of the task was such that participants ended up with the same number of correct trials per condition. However, the total number of trials was variable due to error trials, and was increased on average by 25.1% (SEM=3.2%, N=12). Most error trials were due to movement execution errors (16.3% of all trials, SEM=1.9%, N=12), with that percentage not being significantly affected by cue reliability, cue validity or their interaction (F(2,55)=0.079, p=0.924; F(1,55)=0.728, p=0.397; and F(1,55)=0.127, p=0.723, respectively). During the cue period, the percentage of eye fixation errors (7.1% of all trials, SEM=2.2%, N=12) was not significantly different across cue reliability conditions (F(2,33)=0.241, p=0.787). In contrast, the percentage of premature responses, shown in Figure 1C, increased significantly with cue reliability (F(2,33)=3.923, p=0.030).

### Neural activity

#### Effect of cue reliability

For each channel, we computed the correlation between the log-transformed power of of each frequency band (i.e., delta, theta, alpha, beta, and gamma) and cue reliability during three epochs of interest (i.e., baseline, early cue period, and late cue period). We found a significant cluster of negative correlation (cluster-based p=0.0028) only during the late cue period between the log-transformed power of the beta-band and cue reliability. The power of the beta-band over left central channels that is, contralateral to the responding hand, decreased when reliability increased. We found no significant effect of cue reliability on the power of the beta-band during the baseline period, or the early cue period. In addition, there was no effect of cue reliability on the power of the delta, theta, alpha, and gamma bands in any of the epochs of the task. The map of *t*-values of the correlation between the log-transformed beta power and cue reliability per channel is displayed in Fig. 2A. The channels selected by the cluster-based analysis are identified by the larger symbols. The average cue-aligned time-series of log-transformed beta power from the selected channels is plotted for each cue reliability conditions in Figure 2B, and shows that beta power during the late cue period was scaled negatively with cue reliability.

**Figure 2.**
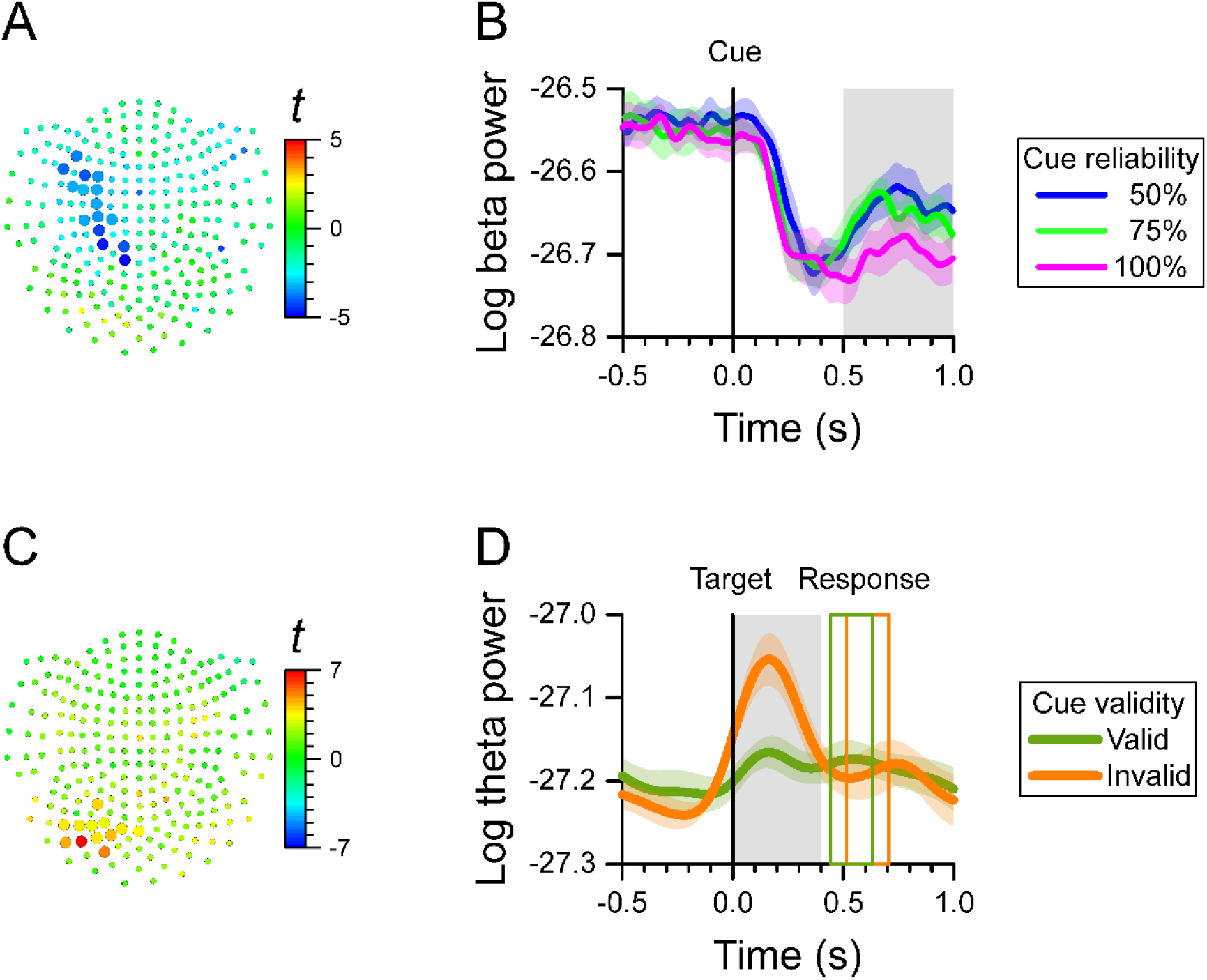
MEG results at the channel level. We found a significant effect of cue reliability on beta power during the late cue period (0.5-1.0 s after cue onset), and a significant cue validity effect on theta power during the reaction time period (0.0-0.4 s after target onset). ***A***, Map of *t*-values for the correlation between log-transformed beta power and cue reliability. Channels selected by the cluster-based analysis are plotted as larger symbols. The negative *t*-values indicate that the greater the cue reliability, the more beta power over left central channels decreased. ***B***, Cue-aligned time-series of log-transformed beta power of the channels selected in ***A*** for the 50%, 75%, and 100% cue reliability conditions. The colored shaded areas indicate the SEM (N=12 participants) for within-subject designs. The gray shaded area shows the time-window of interest. ***C***, Map of *t*-values for the difference in log-transformed theta power between cue validity conditions. Channels selected by the cluster-based analysis are plotted as larger symbols. The positive *t*-values indicate that theta power over left posterior channels was greater for the invalid-cue trials than for the valid-cue trials. ***D***, Target-aligned time-series of log-transformed theta power of the channels selected in ***C*** for the invalid-cue and valid-cue conditions. The colored shaded areas indicate the SEM (N=12 participants) for within-subject designs. The gray shaded area shows the time-window of interest. The colored rectangles show the average onset and offset of the motor response for the valid-cue and invalid-cue conditions.

#### Effect of cue validity

Since cue validity was defined only after target onset, we tested the effect of cue validity on the log-transformed power of each frequency band only during the reaction time period. We found a significant cluster with cue validity effect on theta-band power over left posterior channels (cluster-based p=0.0068), where theta power was greater for invalid-cue trials than for valid-cue trials. We found no significant effect of cue validity on the power of the other bands. The map of *t*-values of the difference in theta power with cue validity, and the channels selected by the cluster-based analysis are displayed in Figure 2C. The average target-aligned time-series of log-transformed theta power from the selected channels is plotted for the two cue validity conditions in Figure 2D, which shows that theta power during the reaction time period was greater for the invalid-cue than for the valid-cue condition.

#### Source analysis of cue reliability and validity effects

The channel-level analysis indicated that there was a significant effect of cue reliability on beta power during the late cue period, and a significant effect of cue validity on theta power during the reaction time period. We performed a source analysis of these effects using the DICS beamforming method. Figure 3 shows that the correlation of log-transformed beta power with cue reliability was primarily localized on the left mid and anterior cingulate cortex (left anterior cingulate included the voxel with the largest absolute *t*-value), the left supplementary motor cortex, the left inferior parietal lobule, the left middle and superior frontal lobules, the left and right striatum (left striatum included the voxel with the second largest absolute *t*-value), and the left and right insula. Figure 4 shows that there was a higher level of theta power for invalid-cue trials than for valid-cue trials primarily on left occipito-parietal areas, including the left precuneus and cuneus (left cuneus included the highest *t*-value), the left superior and middle occipital gyri (left middle occipital gyrus included the second highest *t*-value), the left inferior and superior parietal lobules, and the left mid and posterior cingulate cortex.

**Figure 3.**
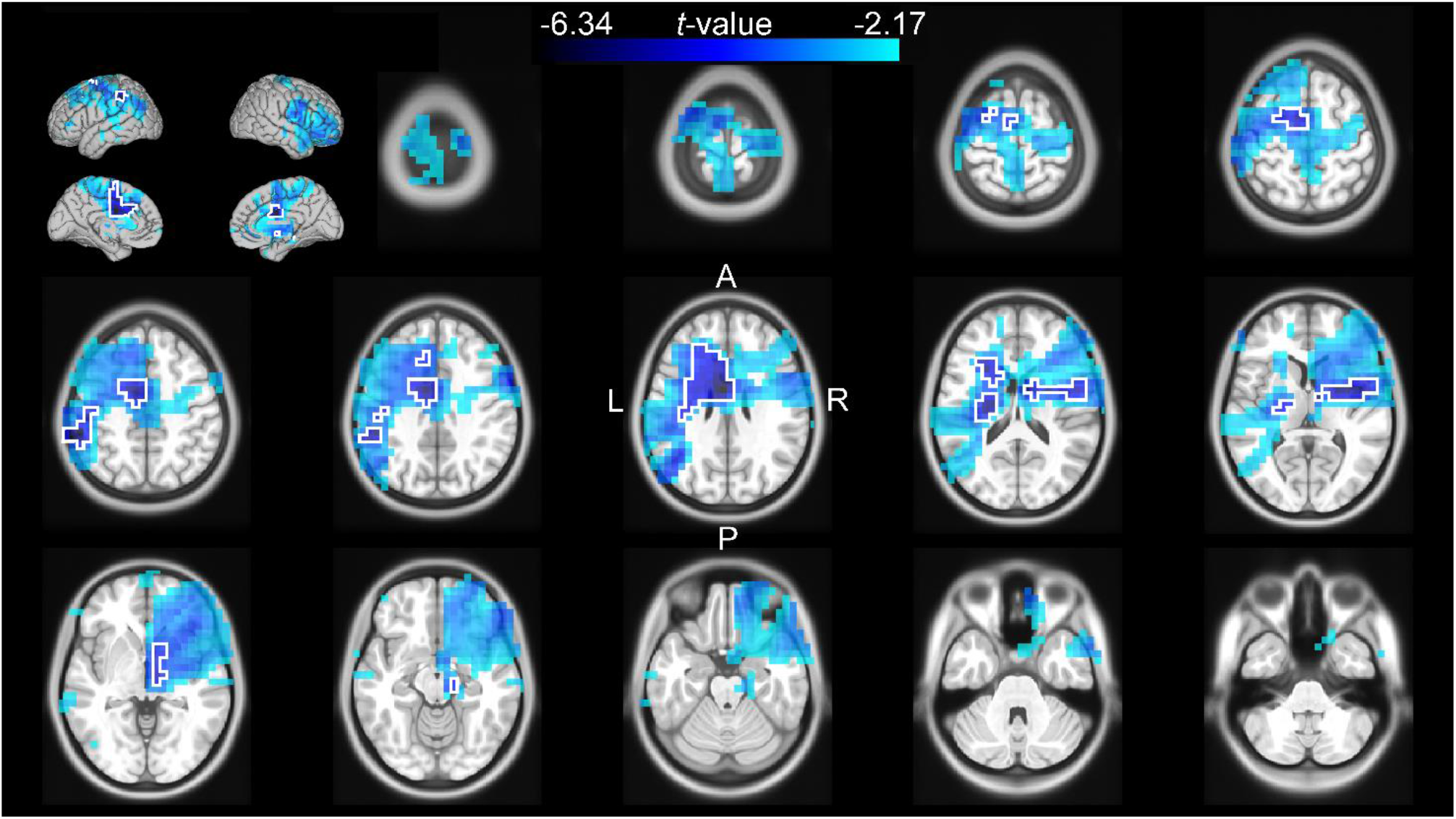
Source analysis of the correlation between the log-transformed beta power and cue reliability during the late cue period (0.5-1s after cue onset). The *t*-values of the correlation are mapped onto the transverse slices and surface of the MNI standard brain. The color scale represents negative *t*-values from the 75^th^ to 100^th^ percentile of the absolute *t*-value distribution. The cluster-based selected area is surrounded by a white border.

**Figure 4.**
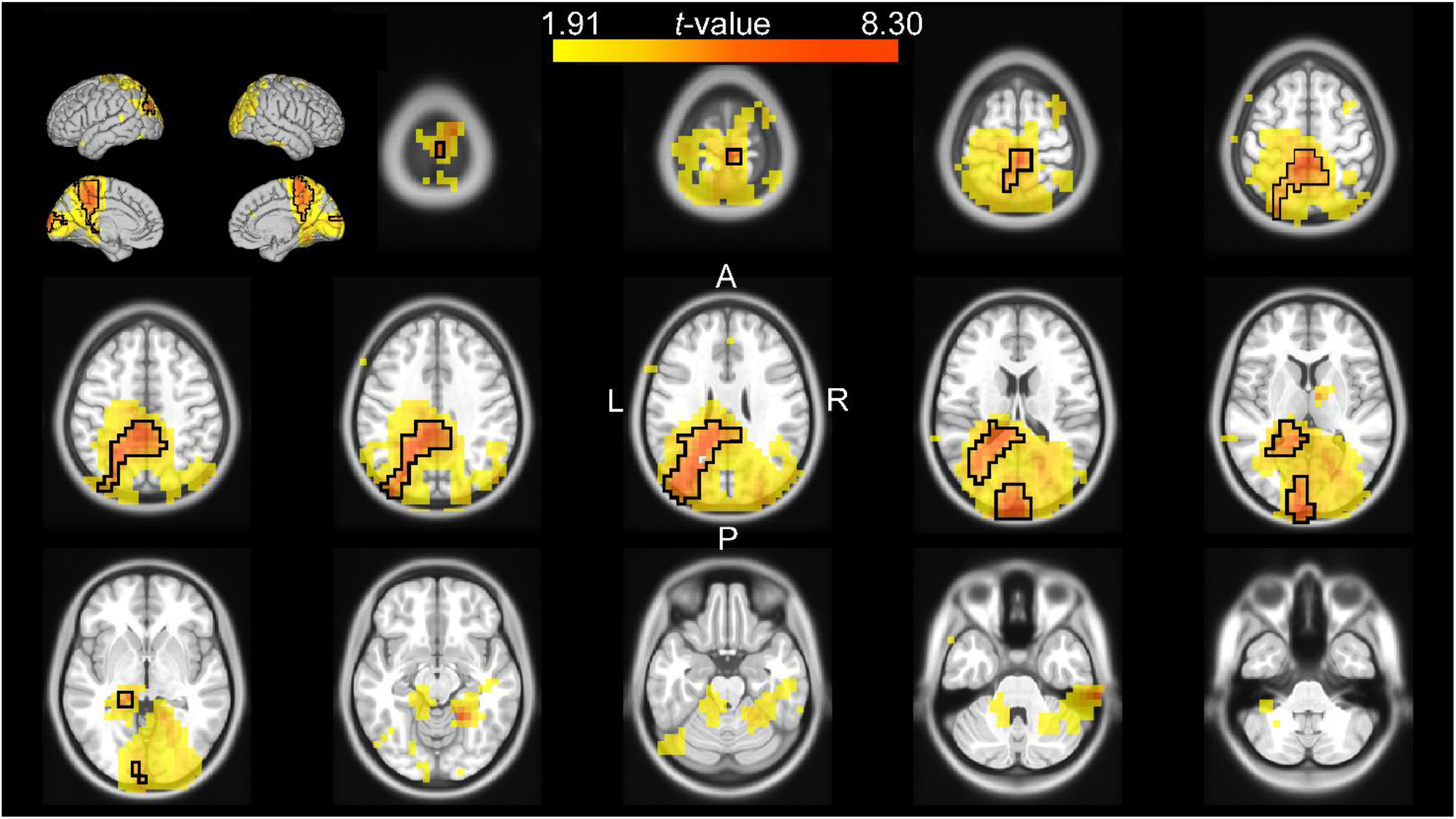
Source analysis of the difference in theta power between valid and invalid cue trials during the reaction time (0-0.4s after target onset). The *t*-values of the difference are mapped onto the transverse slices and surface of the MNI standard brain. The color scale represents positive *t*-values from the 75^th^ to 100^th^ percentile of the absolute *t*-value distribution. The cluster-based selected area is surrounded by a black border.

#### Theta-beta cross-frequency coupling during the reaction time

Given the effect of cue validity on the power of the left occipito-parietal theta-band during the reaction time (Fig. 4), and the well-known decrease in the power of the motor cortical beta-band during the same period, we tested for any cross-frequency coupling between these two brain areas during the reaction time. For this reason, we verified that the main source of decrease of beta power during the reaction time was in the precentral lobule and used voxels in that area for this analysis. Figure 5A shows the left precentral (blue) and occipitoparietal (yellow) voxels used to calculate the theta-beta phase-amplitude PPC, as well as the voxels selected by the cluster-based analysis (red; cluster-based p=0.043) that had a difference in PPC between valid-cue and invalid-cue trials. These voxels were mainly localized in the left mid-occipital cortex. Figure 5B shows the time-course of the PPC in the selected voxels for the valid-cue and invalid-cue trials. Figure 5C depicts the average theta-beta phase-amplitude PPC during the 0.1-0.3 s period after target onset across these voxels for the two cue validity conditions. The presentation of targets that were spatially incongruent from the cue (i.e., invalid cue) was associated with higher PPC values than the presentation of targets spatially congruent with the cue.

We checked whether the difference in PPC between valid-cue and invalid-cue trials described above could be an artefact originating from the difference in theta power also found in these two conditions (Fig. 2D). To this end, we created a bootstrap distribution of difference in PPC between the valid-cue and invalid-cue conditions (N=1,200 samples), using the same target-aligned time-series of theta-band phase used in the analysis above, whereas using time-series of beta-band amplitude taken at a random position in time in each trial. We found that the experimental difference in PPC between valid-cue and invalid-cue trials had only a probability of p=0.0025 to result from the difference in theta power.

**Figure 5.**
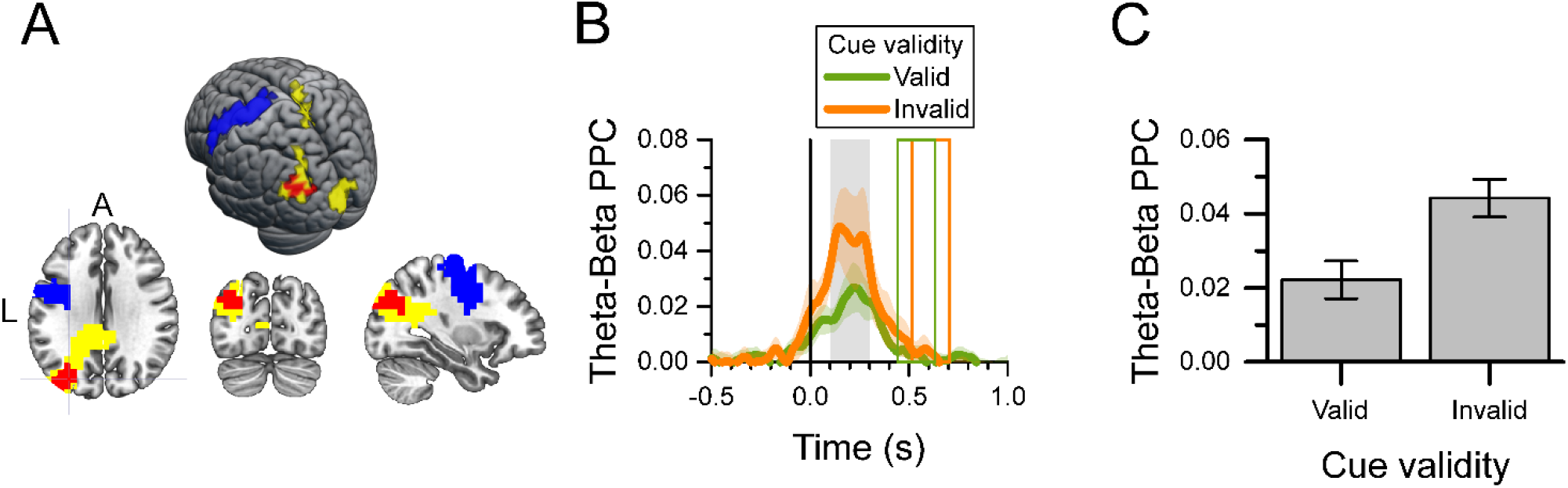
Theta-beta cross-frequency analysis of valid-cue and invalid-cue trials during the 0.1-0.3 s after target onset. ***A***, Map and surface projection on the MNI brain of voxels included in the analysis. The yellow voxels identify those with a significant effect of cue validity (see Fig. 4), and used as the source of the theta-band signal. The blue voxels identify the left precentral gyrus (M1) area used as the source of the beta-band signal. The significant cluster of voxels with a difference in PPC between valid and invalid cue trials selected by the cluster-based analysis are shown in red. ***B***, Target-aligned time-series of theta-beta phase-amplitude PPC per cue validity condition for the red voxels in ***A***. The color shaded area indicates the SEM (N=12) for within-subject design. The grey area identifies the period of interest in the PPC analysis. The colored rectangles show the average onset and offset of the motor response for the valid-cue and invalid-cue conditions. ***C***, Average theta-beta phase-amplitude PPC during the 0.1-0.3 s after target onset (gray rectangle in ***B***) for the selected voxels (red in ***A***) per cue-validity condition. Error bars indicate the SEM (N=12) for within-subject design.

Consequently, it is highly unlikely that the effect of cue validity on PPC was the result of the difference in theta power between the two validity conditions

#### Eye movements after target onset

Since eye movements can affect the power spectrum of MEG signals, we checked whether these could have affected the results relative to cue validity presented above. We found that eye movements outside of the central fixation zone occurred on average across participants in 49.7% (SEM=7.3%) of the trials. There was no significant difference in the percent of trials with eye movements across cue validity conditions (F(1,58)=0.328, p=0.569). When eye movements out of the center occurred, their onset time was significantly shorter for valid-cue than for invalid-cue trials (F(1,52)=11.020, p=0.002). The average eye movement onset was 398 ms (SEM=25 ms) for valid-cue trials and 458 ms (SEM=31 ms) for invalid-cue trials. Note that when there was an eye movement during the reaction time period, its onset time occurred more than 200 ms after the peaks of theta power and PPC described above.

## Discussion

We investigated how the reliability of visual information about the location of the upcoming target modulates the brain mechanisms of motor preparation. To this end, we recorded whole-head MEG signals during a delayed reaching task in which a visual cue provided information about the location of the upcoming target with 50, 75 or 100% reliability. The validity of the cued location was determined by whether the target was presented at the location of the cue or not. In either case, the correct response was to move a joystick-controlled cursor onto the target.

### Reliability of information

We analyzed the change in power of the delta, theta, alpha, beta and gamma frequency bands as a function of cue reliability during the baseline, early cue, and late cue periods of the task. Since cue reliability was determined by the percent of trials in which the cue provided valid information about the upcoming target over a block of trials, we checked whether there was a change in baseline activity that would reflect a change in arousal level between conditions. However, we found no significant effect of reliability condition on baseline activity. In addition, we found no significant effect of reliability condition during the early cue period. However, we found that the power of the beta-band during the late cue period was linearly correlated with the reliability of the information provided by the cue. More specifically, the greater the reliability of the cue, the more the power of the beta-band decreased relative to baseline. This modulation of beta-band power may actually reflect a modulation of the probability of beta-band bursts at the single trial level (Sherman et al., 2016; Little et al., 2019). This effect of cue reliability on the beta-band was observed mostly over left central channels (contralateral to the hand used in the task), and was source-localized to a large network of spatio-motor-related areas, that included the left mid and anterior cingulate cortex, the left and right striatum, the left inferior parietal lobule, the left and right insula, the left supplementary motor cortex, and the left premotor cortex. The source localization of the reliability effect is consistent with findings in other studies of motor attention and decision-making. For example, predictive processing has been linked to activity of a fronto-parietal network that includes prominently the anterior cingulate, striatum, and insula (Ernst and Paulus, 2005; Krain et al., 2006; Balleine et al., 2007; Kuhns et al., 2017; Siman-Tov et al., 2019; Poudel et al., 2020). In addition, pathological conditions that affect decision-making, such as the increased choice impulsivity in Parkinson’s disease (Nombela et al., 2014), have been associated with functional and neuroanatomical abnormalities of the anterior cingulate and striatum (Gescheidt et al., 2013; Ruitenberg et al., 2018; Hlavatá et al., 2019; Kim and Im, 2019). Moreover, neuroimaging, brain-lesion, and stimulation studies have provided evidence for the association of the left inferior parietal cortex with covert attention to an upcoming movement (Rushworth et al., 2001a; Rushworth et al., 2001b; Rushworth et al., 2003). Finally, the supplementary motor area and the premotor cortex, as well as the cingulate cortex are elements of a network that subserves predictive processing (Siman-Tov et al., 2019) and is implicated in the selection and control of motor responses (Ikeda et al., 1999; Dum and Strick, 2002; Wunderlich et al., 2009). Consequently, there is a large fronto-parietal network that is involved in the evaluation of the probability of an upcoming motor response.

We have investigated in previous studies the effect of spatial uncertainty about the location of the upcoming target by varying the number of locations (Pellizzer and Hedges, 2003; Tzagarakis et al., 2010; Tzagarakis et al., 2019), or the size of the sector of space (Pellizzer and Hedges, 2004; Tzagarakis et al., 2015) where the upcoming target could be presented. We have shown that the power of the peri-Rolandic beta-band during motor preparation decreased more, the less uncertain the location of the upcoming target was (Tzagarakis et al., 2010; Tzagarakis et al., 2015; Tzagarakis et al., 2019). However, in those studies spatial uncertainty was conveyed by different displays of visual information (i.e., number of cues or cue size), whereas in the current study the visual display remained the same across cue reliability conditions. Consequently, any difference in motor planning must have been based on the endogenous representation of reliability of information rather than on the exogenous cue. Therefore, the current results corroborate the idea that the change of beta power during motor preparation is associated with the amount of information relative to the upcoming target and not with a concomitant difference in visual display. These results also concur with studies showing that sensorimotor beta power reflects the probabilistic evidence for one motor response over another (Donner et al., 2009; Gould et al., 2012).

Consistent with studies of spatial attention (Jonides, 1980; Posner et al., 1980; Eriksen and Yeh, 1985; Risko and Stolz, 2010; Arjona et al., 2016; Valakos et al., 2020), we found that the reaction time of the reaching response to validly cued trials decreased as the reliability of the cue increased. The effect of cue reliability on motor preparation can also be inferred from the progressive increase of the number of premature response errors as cue reliability increased. These results can be explained by the effect of cue reliability on beta-power during the cue period. The lower the level of beta-band power during the late cue period, the closer it is to the level reached during the motor response, hence the shorter reaction time and the greater likelihood of a premature movement onset (Tzagarakis et al., 2010; Tzagarakis et al., 2015). This is also consistent with the finding that people with high impulsivity scores have a greater decrease of beta-band power during motor preparation and a higher number of premature responses than people with low impulsivity scores (Tzagarakis et al., 2019; Barth et al., 2021).

### Validity of information

We analyzed the power of the delta, theta, alpha, beta, and gamma frequency bands after target onset, that is, once the validity status of the cue was determined. This analysis showed that there was a significantly greater phasic increase of theta-band power after target onset for invalid-cue trials than for valid-cue trials. This effect was observed mainly over left posterior channels, and was source-localized to the left occipito-parietal cortex and the left mid and posterior cingulate. The source localization is consistent with the role of the occipito-parietal region in visual attention (Wojciulik and Kanwisher, 1999; Shomstein, 2012) and with the role of the posterior cingulate in regulating the focus of attention (Leech and Sharp, 2014). Furthermore, the left posterior parietal region is a key component of motor attention (Rushworth et al., 2003), whereas the left middle occipital gyrus plays a critical role in the proper regulation of attention (Proal et al., 2011; Sörös et al., 2017). The effect of cue validity on the power of theta is consistent with other studies using cueing tasks that have shown greater theta power and greater amplitude of event-related potentials for invalid-cue than for valid-cue trials (Rawle et al., 2012; Arjona et al., 2016; Proskovec et al., 2018; Valakos et al., 2020). There is actually a strong relation between the effect of cue validity on theta power and the P300 event-related potential (Yordanova and Kolev, 1998), and both are thought to reflect the same phenomenon (Başar-Eroglu et al., 1992; Wang and Ding, 2011). Note that there was a phasic increase of power of theta also after cue onset for the same channels (data not shown). For these reasons, the phasic increase of theta power seems to signal the onset of an unexpected stimulus and the need to shift attention. These results are also consistent with studies that found transient change of hemodynamic response in the posterior parietal cortex associated with shift of spatial attention (Corbetta et al., 2000; Yantis et al., 2002). Finally, the absence of an effect on beta power after target onset concurs with previous findings showing that beta-band power reaches the same level during motor execution regardless of its previous level during the preparatory period (Tzagarakis et al., 2010; Grent-’t-Jong et al., 2015; Heinrichs-Graham and Wilson, 2016).

In order to further elucidate the integration of attentional modulation and motor control, we analyzed the theta-beta cross-frequency coupling during the reaction time. Given the close correspondence between the increase in theta power and event-related potentials (Başar-Eroglu et al., 1992; Wang and Ding, 2011), the theta-beta coupling could also be interpreted as a coupling between visuo-spatial event-related potentials and motor-related beta oscillations. More extensive coupling analyses could be performed using other frequency bands and brain regions. For example, theta-gamma coupling has been considered as a mechanism for the coding spatial information in memory (Lisman and Jensen, 2013). However, here we opted for an approach that restricted the probability of type I error and purposely limited the analysis to the frequency bands and brain regions identified in the previous analyses. Furthermore, analyses of functional connectivity are known to be susceptible to spurious results if careful precautions are not taken (Palva and Palva, 2012; Aru et al., 2015; Bastos and Schoffelen, 2015). To mitigate the effect of field-spread of activity across sensors, we performed the connectivity analysis in source space. The LCMV beamformer estimates the activity of a source while suppressing the contributions of all the other sources (Van Veen et al., 1997), which is why it is appropriate for connectivity analysis unlike sensor-based signals (Schoffelen and Gross, 2009; Palva and Palva, 2012). The results showed greater phasic theta-beta cross-frequency coupling between the left middle occipital cortex and the left motor cortex during invalid-cue trials than during valid-cue trials. We showed that the effect of cue validity on theta-beta coupling was not an artefact occurring from the difference in theta-band power between validity conditions (Schoffelen and Gross, 2009; Aru et al., 2015; Bastos and Schoffelen, 2015). Consequently, this result suggests that theta-band activity provided a visuo-spatial signal to the motor region signaling that a change of the motor response was necessary. This updating may explain the longer reaction time for invalid-cue trials than for valid-cue trials. However, it is not clear at this point whether the cross-frequency coupling just signals that an update of the planned motor response is necessary, or whether it carries more specific information about what the required update should be. Further investigations will be needed to understand better the mechanisms by which the motor plan is updated and the role of the cross-frequency coupling in this process.

Finally, as regards the brain areas involved in these mechanisms, it is generally considered that visuo-spatial orienting is dependent on a fronto-parietal network predominantly lateralized to the right hemisphere (Corbetta and Shulman, 2002). However, the attention requirement of targeted-limb movements is associated with the left posterior parietal cortex (Rushworth et al., 2001a; Rushworth et al., 2001b), in contrast to visuospatial detection or discrimination tasks used in typical studies of orienting and shifting attention (Corbetta and Shulman, 2002). The absence of a significant modulation of frontal theta-band activity with cue validity in the current study may reflect the weaker and less consistent involvement of frontal regions than posterior parietal regions in attention shifting (Wojciulik and Kanwisher, 1999; Wager et al., 2004). Furthermore, the modulation of frontal theta-band activity was found to be associated with response inhibition (Yamanaka and Yamamoto, 2010; Isabella et al., 2015). For these reasons, the effect of cue validity over the occipito-parietal cortex may reflect the more preponderant role of re-orienting attention in our task than in go/no-go tasks.

### Limitations

This study has several limitations. The task had only three different levels of cue reliability, which restricted the quantitative description of the relation between beta power and cue reliability. We analyzed the linear relation between the log-transformed beta power and the percent of valid cues in a block of trials, and found that there was a significant linear component in the progression between the decrease in beta power and the increase in cue reliability. This does not exclude that the relation between these two variables is more complex than the linear relation tested here. However, more levels of cue reliability would be needed to perform a detailed analysis of the mathematical function that best describes their relation. Another limitation resulted from keeping the initial cue on the display throughout the trial regardless of whether the target was congruent with it or not. This means that invalid-cue trials were visually different than valid-cue trials which raises the possibility that any neural effect of cue validity was confounded with the neural effect of a different visual stimulus. However, since the effect of cue validity on the theta-band was consistent with the results of other studies in which a visual stimulus appeared at an unexpected location (Rawle et al., 2012; Arjona et al., 2016; Proskovec et al., 2018), we believe that the difference in visual display was not the main factor in the occurrence of this effect. In addition, the phase-amplitude effect of mid-occipital theta activity on beta oscillations in the motor cortex suggests that the task elicited mechanisms that went beyond perceptual processing and directly affected motor planning. Nevertheless, further work is needed to precisely disentangle the effects of visual response and the motor information carried by it.

### Conclusions

We have examined neural oscillatory activity associated with cue reliability and validity in a delayed reaching task. During the cue period, the power of the beta-band over a large network of motor-related areas decreased as cue reliability increased. The greater the decrease in beta-band power, the closer the motor system was to execute the response, which shortened the reaction time and increased the chances of a premature response. After target onset, the power of the theta-band at left occipito-parietal areas had a greater phasic increase during invalid-cue trials than during valid-cue trials. In addition, there was a greater theta-beta cross-frequency coupling after target onset between the middle occipital cortex and the motor cortex during invalid-cue trials than during valid-cue trials. Consequently, theta-band activity seemed to have signaled the need to reorient visuo-spatial attention and to change motor plan. The change in motor plan was associated with an increase in reaction time. In summary, beta-band power in several motor-related areas reflected the reliability of directional information used during motor preparation, whereas theta-band power of occipito-parietal areas signaled whether that information was valid or not. These results elucidate mechanisms of interaction between visuo-spatial attention and motor processes.

## Acknowledgements

The study was supported by the U.S. Department of Veterans Affairs Clinical Sciences Research and Development Merit Review Awards I01 CX000437 and I01 CX001773, and by the United States Department of Veterans Affairs Rehabilitation Research and Development Pilot Project I21 RX003007. We thank Arthur Leuthold, PhD and Dale Boeff from the Minneapolis Veterans Affairs Health Care System for their expert technical assistance.

